# Longitudinal Epidemiological Study of Autism Subgroups Using Autism Treatment Evaluation Checklist (ATEC) Score

**DOI:** 10.1101/266221

**Authors:** Shreyas Mahapatra, Edward Khokhlovich, Samantha Martinez, Benjamin Kannel, Stephen M Edelson, Andrey Vyshedskiy

## Abstract

Here we report the results of the subgroup analyses of an observational cohort of children whose parents completed the Autism Treatment Evaluation Checklist (ATEC) over the period of several years. A linear mixed effects model was used to evaluate longitudinal changes in ATEC scores within different patient subgroups. All groups decreased their mean ATEC score over time indicating improvement of symptoms, however there were significant differences between the groups. Younger children improved more than the older children. Children with milder ASD improved more than children with more severe ASD in the Communication subscale. There was no difference in improvement between females vs. males. One surprising finding was that children from developed English-speaking countries improved less than children from non-English-speaking countries.

## Introduction

Design considerations for an ASD early-intervention clinical trial must take into account (1) the trial duration, (2) number of participants, and (3) the quality of participant assessment. A short clinical trial of an early therapeutic intervention in 2 to 3 year old children can easily miss a target, as an improvement of symptoms may not emerge until children reach the school age. Small numbers of participants can easily skew the data as ASD is known to be a highly heterogeneous disorder. Longer clinical trials, with a greater number of participants, provide a better test for any intervention. Increasing the trial duration and the number of trial participants, however, raises the demand for regular assessment of participants by trained psychometric technicians. Furthermore, to attain the larger number of trial participants, clinical trials must accept participants across a large geographical region. The logistical issues associated with such an endeavor come at immense cost. As a result, large numbers of ASD clinical trials working under a limited budget suffer from short duration and low participant number, often compromising the trial objectives (e.g., (Drew et al., 2002; Whitehouse et al., 2017).

A parent-completed Autism Treatment Evaluation Checklist (ATEC) assessment tool was in part designed to circumvent these problems (Rimland & Edelson, 1999). If caregivers could serve as psychometric technicians and conduct regular evaluations of their children, the cost of clinical trials would be substantially reduced while simultaneously allowing for longer trial duration. There is an understanding in the psychological community that parents cannot be trusted with an evaluation of their own children. In fact, parents often yield to wishful thinking and overestimate their children’s abilities on a single assessment. However, the pattern of changes can be generated by measuring the score dynamics over multiple assessments. When a single parent completes the same evaluation every three months over multiple years, changes in the score become meaningful. ATEC was specifically designed to measure changes in ASD severity, making it useful in monitoring behaviors over time as well as tracking the efficacy of a treatment. ATEC is comprised of four subscales: 1) Speech/Language/Communication, 2) Sociability, 3) Sensory/Cognitive Awareness, and 4) Health/Physical/Behavior. The subscales provide survey takers with the information about specific areas of behaviors which may change over time.

The current observational study was initiated nearly two decades ago when one of the authors (Stephen M. Edelson of Autism Research Institute) distributed ATEC questionnaire to parents of children with ASD. Initially, ATEC evaluations were distributed as hard copy. In 2013 the online version of ATEC was developed. The current study analyzed data reported by participants using the online version ATEC over a four-year time span (2013 to 2017). The goal of the study was to characterize the typical changes in ATEC score over time as a function of children age, sex, ASD severity, and country of origin in a large and diverse group of participants.

## Methods

### ATEC Evaluation Structure

The ATEC is a caregiver-administered questionnaire designed to measure changes in severity of ASD in response to treatment. A total score and four subscale scores are reported. Questions in the first three subscales are scored using a 0-2 scale. The fourth subscale, Health/Physical/Behavior, is scored using a 0-3 point scale. ATEC can be accessed online or in hard-copy format.

The first subscale, Speech/Language/Communication, contains 14 items and its score ranges from 0 to 28 points. The Sociability subscale contains 20 items within 0 to 40 score range. The third subscale, Sensory/Cognitive awareness, has 18 items and scores range from 0 to 36. Finally, the Health/Physical/Behavior subscale contains 25 items. The scores from each subscale are combined in order to calculate a Total Score, which ranges from 0 to 179 points. A lower score indicates a lower severity of ASD symptoms and a higher score correlates with more severe symptoms of ASD.

### Collection of Evaluations

ATEC responses were collected from participants voluntarily completing online ATEC evaluations from 2013 to 2017. The ATEC questionnaire was not actively advertised and use primarily originated from online searches. Participants consented to anonymized data analysis and publication of the results.

### Evaluations of ATEC score changes over time

In order to study how ATEC scores change overtime and whether those changes vary within different ASD subgroups, the concept of a "Visit" was developed by dividing the two-year-long observation interval into 3-month periods. All evaluations were mapped into 3-month-long bins with the first evaluation placed in the first bin. When more than one evaluation was completed within a bin, their results were averaged to calculate a single number representing this 3-month interval. It was then hypothesized that there was an interaction between a Visit and a given subgroup category (age, sex, ASD severity, and country of origin). Statistically, this hypothesis was modeled by applying Linear Mixed Effect (LME) model with repeated measures, where an interaction term was introduced to test the hypothesis. This, in turn, enabled generation of pairwise differences between modeled subgroups at different visits. Participant specific variability was accounted for by introducing random effect into the model.

Pairwise differences were computed by applying “LS Means “ and “contrast” functions to a generated LME model. For each ATEC score and a given subgroup, the following output was generated:

1. General ANOVA summary of the model itself, including p-values for each covariate and the interaction term among them
2. LS Means computed for a given category at each visit (with 95% Confidence Interval)
3. Pairwise differences between categories at different visits with p-values adjusted for multiple comparisons testing using Tukey method.

## Participants

Participants were selected based on the following criteria:

1) **Completeness:** Participants who did not provide a date of birth (DOB) were excluded. As participants’ DOB were utilized to determine age, the availability of DOB was a necessary.

2) **Consistency:** Participants had to have completed at least three questionnaires within 2 years and the interval between the first and the last evaluation was one year or longer.

3) **Maximum age:** Participants older than twelve years of age were excluded from this study.

As diagnosis was not part of ATEC questionnaire, some neurotypical participants could be present in the database. To limit the contribution from neurotypical children, we excluded participants that may have represented the neurotypical population by using the *Minimum age* and *the Minimal ATEC severity* criteria.

4) **Minimum age:** Participants who completed their first evaluation before the age of 2 were excluded from this study, as the diagnosing of ASD in this age group is uncertain and the parents of some of these young cases may have completed the ATEC because they wanted to check whether their normal child had signs of autism.

5) **Minimal ATEC severity:** Participants with initial ATEC scores of less than 20 were excluded.

After excluding participants that did not meet these criteria, there were 2272 total participants.

### Age groups

Participants were grouped based on age, calculated from the date of birth at the time of the first completed evaluation. The three age groups were: 2-3 years of age (YOA), 3.1-6 YOA, 6.1-12 YOA (Table 1).

**Table 1.** Characteristics and baseline measures for age groups.

| Table 1 |  |  |  |  |
| --- | --- | --- | --- | --- |
| *Characteristics and baseline measures for age groups.* |  |  |  |  |
| Age (abbreviation used in the paper) | Participants in each age group (total) | Participants in each age group (%) | Age at baseline (mean ± SD) | Initial ATEC total score (mean ± SD) |
| 2-3 YOA (2-3) | 407 | 18 | 2.58±0.30 | 71±22 |
| 3.1-6 YOA (3-6) | 1205 | 53 | 4.34±0.83 | 60±24 |
| 6.1-12 YOA (6-12) | 660 | 29 | 8.19±1.61 | 61±24 |

### Autism Severity Measurements

The initial ATEC total score was used as proxy for ASD severity. Participants were organized into three groups: mild (initial ATEC total score 20-49), moderate (initial ATEC total score 50-79), and severe (initial ATEC total score > 80) (Table 2).

**Table 2.** Characteristics and baseline measures for ASD severity groups

| Table 2 |  |  |  |  |  |
| --- | --- | --- | --- | --- | --- |
| *Characteristics and baseline measures for ASD severity groups* |  |  |  |  |  |
| Autism Severity | Initial ATEC Total Score | Participants in each severity group (total) | Participants in each severity group (%) | Age at baseline (mean ± SD) | Initial ATEC total score (mean ± SD) |
| Mild | 20-49 | 741 | 33 | 5.47±2.35 | 47±8 |
| Moderate | 50-79 | 999 | 44 | 5.06±2.41 | 64±8 |
| Severe | 80 ≤ | 532 | 23 | 5.11±2.59 | 95±14 |

### Country groups

Participants were split into two groups based on their country of origin. The developed English-speaking nations included participants from the United States, Canada, United Kingdom, Ireland, Australia, and New Zealand. Participants from other countries were grouped together as “the non-English-speaking countries” (Table 3). In the non-English-speaking countries group, only 53 participants (4%) were from Japan, France, Germany and northern Europe. The majority of participants were from Latin America (859; 63%), southern Europe (182; 13%), and India (70; 5%).

**Table 3.** Characteristics and baseline measures for the English-speaking countries and the non-English-speaking countries.

| Group | Participants in each group (total) | Participants in each group (%) | Age at baseline (mean ± SD) | Initial ATEC total score (mean ± SD) |
| --- | --- | --- | --- | --- |
| English-speaking countries | 972 | 43 | 5.67±2.56 | 62±24 |
| Non-English-speaking countries | 1294 | 57 | 4.85±2.28 | 63±24 |

### Sex groups

Data were stratified based on sex. 83% of the 2272 participants were males (Table 4).

**Table 4.**
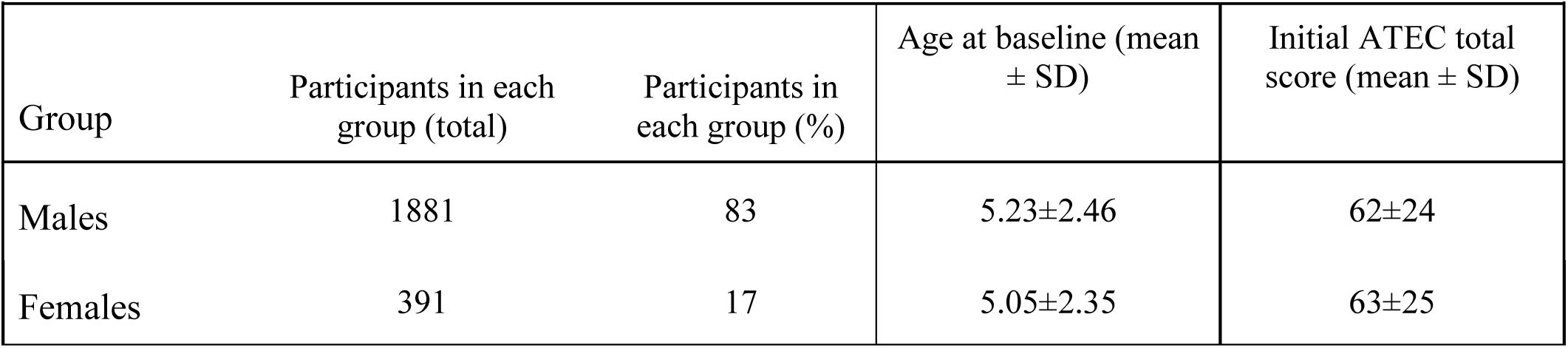
Characteristics and baseline measures for gender groups

## Results

The least squared means (LS Means), as well as the dynamics of score changes (LS Means differences) over time (between visits) for each group are presented in the supplementary materials. The fitting of LME model allowed us not only to assess the temporal dynamics of the scores, but also to evaluate the “tightness” of each individual mean value by generating 95% confidence interval. There was a high degree of data consistency, similar to what was reported by Magiati (Magiati, Moss, Yates, Charman, & Howlin, 2011). The differences between participant subgroups at different visits, and differences between the first and the last visit per subgroup are discussed in greater detail below.

### Effect of sex on longitudinal change of ATEC scores

The interaction term between Sex group and Visits was not statistically significant (at the α = 0.05 level of significance) for either the ATEC total score or any of the subscale scores (Table S1).

### Longitudinal change of ATEC scores as a function of age

The significance of interaction term between Age group and Visits shows that the dynamics of ATEC total score as well as scores in the Communication, Sociability, and Sensory subscales vary within different Age groups (Table S2). Table 5 shows LS Means for age groups at the initial and the last visits. Reduction in ATEC total score (showing the degree of improvement) was inversely related to age (Table 5). Over the two years the 2-3 YOA group improved by 28.35 units (SE=1.30, p<0.0001), the 3-6 YOA group improved by 19.73 units (SE=0.72, p<0.0001), and the 6-12 YOA group improved by 13.80 units (SE=0.96, p<0.0001).

**Table 5:** LS Means for various age groups. Data are presented as: LS Mean (SE; 95% CI). The difference between Visit 8 and Visit 1 is presented as LS Mean (SE; P-value).

|  | Visit 1 |  |  | Visit 8 |  |  | Visit 8 – Visit 1 |  |  |
| --- | --- | --- | --- | --- | --- | --- | --- | --- | --- |
|  | 2-3 YOA | 3-6 YOA | 6-12 YOA | 2-3 YOA | 3-6 YOA | 6-12 YOA | 2-3 YOA | 3-6 YOA | 6-12 YOA |
| **ATEC Total** | 64.41<br>(0.76;<br>62.93-65.90) | 62.15<br>(0.47;<br>61.24-63.07) | 61.98<br>(0.61;<br>60.78-63.17) | 36.06<br>(1.28;<br>33.55-38.58) | 42.42<br>(0.72;<br>41.01-43.84) | 48.18<br>(0.95;<br>46.32-50.04) | -28.35<br>(1.30;<br><0.0001) | -19.73<br>(0.72;<br><0.0001) | -13.80<br>(0.96;<br><0.0001) |
| **Subscale 1**<br>**Communication** | 15.58<br>(0.18;<br>15.23-15.93) | 15.12<br>(0.11;<br>14.90-15.33) | 14.84<br>(0.14;<br>14.56-15.12) | 7.07<br>(0.29;<br>6.49-7.65) | 10.42<br>(0.16;<br>10.10-10.74) | 12.82<br>(0.22;<br>12.40-13.25) | -8.51<br>(0.29;<br><0.0001) | -4.70<br>(0.16;<br><0.0001) | -2.02<br>(0.22;<br><0.0001) |
| **Subscale 2**<br>**Sociability** | 13.84<br>(0.23;<br>13.39-14.28) | 13.11<br>(0.14;<br>12.84-13.39) | 13.27<br>(0.18;<br>12.90-13.63) | 6.92<br>(0.40;<br>6.12-7.71) | 8.87<br>(0.23;<br>8.42-9.31) | 10.05<br>(0.30;<br>9.47-10.65) | -6.92<br>(0.42;<br><0.0001) | -4.25<br>(0.23;<br><0.0001) | -3.21<br>(0.31;<br><0.0001) |
| **Subscale 3**<br>**Sensory** | 14.80<br>(0.22;<br>14.37-15.22) | 14.26<br>(0.13;<br>13.99-14.52) | 14.31<br>(0.18;<br>13.96-14.65) | 14.80<br>(0.22;<br>14.37-15.22) | 14.26<br>(0.13;<br>13.99-14.52) | 14.31<br>(0.18;<br>13.96-14.65) | -6.35<br>(0.38;<br><0.0001) | -4.51<br>(0.21;<br><0.0001) | -3.66<br>(0.28;<br><0.0001) |
| **Subscale 4**<br>**Physical** | 19.94<br>(0.34;<br>19.28-20.60) | 20.10<br>(0.21;<br>19.70-20.51) | 20.61<br>(0.28;<br>20.07-21.15) | 13.26<br>(0.59;<br>12.09-14.42) | 13.67<br>(0.33;<br>13.02-14.33) | 15.59<br>(0.44;<br>14.73-16.45) | -8.51<br>(0.20;<br><0.0001) | -4.70<br>(0.16;<br><0.0001) | -2.02<br>(0.22;<br><0.0001) |

For age group comparisons, three pairwise comparisons were made (Table 6). Neither pairwise comparison reached statistical significance at Visit 1, but all three pairwise comparisons in ATEC total score yielded statistically significant differences in LS Means at Visit 8 with younger children improving more than the older children. The difference in ATEC total score for the 2-3 YOA group relative to the 3-6 YOA group was −2.26 units (SE=0.83, p=0.6501) at Visit 1 and −6.36 units (SE=1.44, p=0.0042) at Visit 8. ATEC total score difference for the 2-3 YOA group relative to the 6-12 YOA group was 2.44 units (SE=0.93, p=0.7182) at Visit 1 and −12.12 units (SE=1.57, p<0.0001) at Visit 8. ATEC total score difference for the 3-6 YOA group relative to the 6-12 YOA group was 0.17 units (SE=0.71, p=1.0000) at Visit 1 and - 5.76 units (SE=1.16, p<0.0003) at Visit 8. These observations were recapitulated in the Communication and Physical subscales (Table 6). For the Sociability subscale, only the 2-3 YOA group vs. 3-6 YOA group and 2-3 YOA group vs. 6-12 YOA group yielded a statistically significant decrease in score at Visit 8 (Table 6). For the Sensory subscale, only the 2-3 YOA group vs. 6-12 YOA group yielded a statistically significant decrease in score at Visit 8 (Table 6).

**Table 6:**
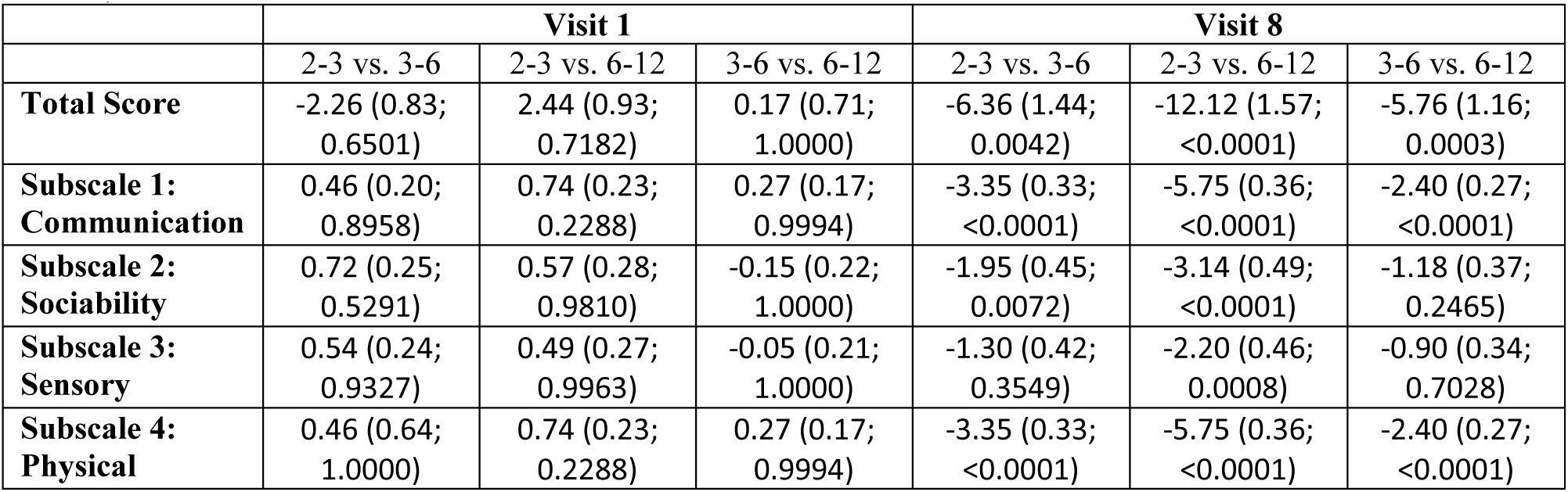
LS Mean differences between Age Groups. Data are presented as: LS Mean difference (SE; P-Value)

### Country effects on ATEC scores

Surprisingly, a comparison of developed English-speaking nations (the United States, Canada, United Kingdom, Ireland, Australia, and New Zealand) to non-English-speaking countries demonstrated greater improvements in ATEC total score and all subscales in the non-English-speaking nations. The significance of interaction term between Country group and Visits shows that the dynamics of ATEC total score and all subscales varies in different Country groups (Table S13). Table 7 shows LS Means for Country groups at the initial and the last visits. Reduction in ATEC total score was greater in non-English-speaking nations group (Table 7). Over the period of two years the participants in the English-speaking nations group improved by 16.70 units (SE=0.80, p<0.0001), and non-English-speaking nations group improved by 21.58 units (SE=0.70, p<0.0001).

**Table 7:** LS Means of English-speaking countries group and non-English-speaking countries group. Data are presented as LS Mean (SE; 95% CI). The difference between Visit 8 and Visit 1 is presented as LS Mean (SE; P-value).

|  | Visit 1 |  | Visit 8 |  | Visit 8 – Visit 1 |  |
| --- | --- | --- | --- | --- | --- | --- |
|  | Non-English-Speaking Countries | English-Speaking Countries | Non-English-Speaking Countries | English-Speaking Countries | Non-English-Speaking Countries | English-Speaking Countries |
| **ATEC Total** | 62.92 (0.67; 61.61-64.23) | 62.09 (0.69; 60.74-63.44) | 41.34 (0.86; 39.66-43.03) | 45.40 (0.93; 43.58-47.21) | -21.58 (0.70; <0.0001) | -16.70 (0.80; <0.0001) |
| **Subscale 1 Communication** | 15.54 (0.16; 15.23-15.86) | 14.98 (0.16; 14.66-15.30) | 10.54 (0.20; 10.14-10.93) | 11.09 (0.22; 10.66-11.52) | -5.01 (0.16; <0.0001) | -3.89 (0.19; <0.0001) |
| **Subscale 2 Sociability** | 13.42 (0.19; 13.04-13.81) | 13.29 (0.20; 12.89-13.69) | 8.35 (0.26; 7.84-8.86) | 9.73 (0.28; 9.17-10.28) | -5.07 (0.22; <0.0001) | -3.57 (0.26; <0.0001) |
| **Subscale 3 Sensory** | 14.33 (0.19; 13.96-14.70) | 14.31 (0.20; 13.92-14.69) | 9.35 (0.25; 8.86-9.83) | 10.24 (0.27; 9.71-10.76) | -4.98 (0.21; <0.0001) | -4.07 (0.24; <0.0001) |
| **Subscale 4 Physical** | 20.06 (0.29; 19.49-20.63) | 20.37 (0.30; 19.78-20.97) | 13.42 (0.38; 12.68-14.19) | 15.02 (0.42; 14.20-15.84) | -6.63 (0.33; <0.0001) | -5.35 (0.38; <0.0001) |

The difference in ATEC total score for the English-Speaking Counties group relative to the non-English-speaking countries was 0.83 units (SE=0.61, p = 0.9937) at Visit 1 and −4.05 units (SE=1.02, p=0.0056) at Visit 8 (Table 8). This statistically significant difference at Visit 8 indicates that children in the English-speaking countries improve their symptoms to a smaller degree than children in the non-English speaking nations. Dissection of the ATEC total score into subscales indicated that children in the non-English speaking nations demonstrated greater improvements of their symptoms in all subscales with the Sociability and Physical subscales having the greatest contribution to the difference between the groups. The difference in the Sociability subscale score for the English-Speaking Counties group relative to the non-English-speaking countries was 0.13 units (SE=0.19, p = 1.0000) at Visit 1 and −1.37 units (SE=0.32, p = 0.0023) at Visit 8 (Table 8). The difference in the Physical subscale score for the English-Speaking Counties group relative to the non-English-speaking countries was −0.32 units (SE=0.28, p = 0.9990) at Visit 1 and −1.60 units (SE=0.47, p = 0.0619) at Visit 8 (Table 8).

**Table 8:** LS Mean differences between the English-speaking countries group and non-English-speaking countries group. Data are presented as LS Mean difference (SE; P-Value)

|  | Visit 1 | Visit 8 |
| --- | --- | --- |
|  | Non-English-Speaking Countries vs. English-Speaking Countries | Non-English-Speaking Countries vs. English-Speaking Countries |
| **Total Score** | 0.83 (0.61; 0.9937) | -4.05 (1.02; 0.0056) |
| **Subscale 1: Communication** | 0.56 (0.15; 0.0115) | -0.55 (0.24; 0.6330) |
| **Subscale 2: Sociability** | 0.13 (0.19; 1.0000) | -1.37 (0.32; 0.0023) |
| **Subscale 3: Sensory** | 0.02 (0.18; 1.0000) | -0.89 (0.30; 0.1845) |
| **Subscale 4: Physical** | -0.32 (0.28; 0.9990) | -1.60 (0.47; 0.0619) |

### Change of ATEC scores as a function of ASD severity

The significance of interaction term between ASD severity group and Visits shows that the dynamics of ATEC total score and of individual scores within all subscales differs between severity groups (Table S24). Table 9 shows the LS Mean calculations for the three severity groups (mild, moderate, severe) at Visit 1 and Visit 8. Reduction in ATEC total score was directly related to severity (Table 9). Over the two years, the mild group improved by 11.20 units (SE=0.87, p<0.0001), the moderate group by 20.56 units (SE=0.76, p<0.0001), and the severe group by 29.52 units (SE=1.10, p<0.0001).

**Table 9:** LS Means for various severity groups. Data are presented as: LS Mean (SE; 95% CI). The difference between Visit 8 and Visit 1 is presented as LS Mean (SE; P-value).

|  | Visit 1 |  |  | Visit 8 |  |  | Visit 8 – Visit 1 |  |  |
| --- | --- | --- | --- | --- | --- | --- | --- | --- | --- |
|  | Mild | Moderate | Severe | Mild | Moderate | Severe | Mild | Moderate | Severe |
| **ATEC Total** | 56.44<br>(0.92;<br>54.64-58.25) | 62.94<br>(0.69;<br>61.60-64.29) | 70.57<br>(1.10;<br>68.40-72.73) | 45.25<br>(1.13;<br>43.02-47.47) | 42.38<br>(0.90;<br>40.61-44.16) | 41.04<br>(1.41;<br>38.28-43.81) | -11.20<br>(0.87;<br><0.0001) | -20.56<br>(0.76;<br><0.0001) | -29.52<br>(1.10;<br><0.0001) |
| **Subscale 1: Communication** | 14.62<br>(0.19;<br>14.25-14.99) | 15.41<br>(0.17;<br>15.09-15.74) | 15.94<br>(0.20;<br>15.55-16.33) | 10.96<br>(0.25;<br>10.49-11.46) | 10.49<br>(0.22;<br>10.07-10.92) | 10.89<br>(0.29;<br>10.31-11.46) | -3.64<br>(0.21;<br><0.0001) | -4.92<br>(0.18;<br><0.0001) | -5.05<br>(0.27;<br><0.0001) |
| **Subscale 2: Sociability** | 10.93<br>(0.23;<br>10.48-11.39) | 13.44<br>(0.20;<br>13.05-13.84) | 16.39<br>(0.26;<br>15.88-16.91) | 9.07<br>(0.31;<br>8.45-9.69) | 8.61<br>(0.28;<br>8.07-9.15) | 9.01 (0.39;<br>8.25-9.76) | -1.87<br>(0.28;<br><0.0001) | -4.83<br>(0.25;<br><0.0001) | -7.39<br>(0.36;<br><0.0001) |
| **Subscale 3: Sensory** | 12.70<br>(0.23;<br>12.26-13.15) | 14.48<br>(0.20;<br>14.09-14.87) | 16.22<br>(0.25;<br>15.72-16.72) | 10.09<br>(0.30;<br>9.50-10.68) | 9.59<br>(0.26;<br>9.07-10.11) | 9.30 (0.37;<br>8.58-10.02) | -2.62<br>(0.26;<br><0.0001) | -4.89<br>(0.23;<br><0.0001) | -6.92<br>(0.33;<br><0.0001) |
| **Subscale 4: Physical** | 17.51<br>(0.34;<br>16.85-18.17) | 19.95<br>(0.30;<br>19.36-20.54) | 24.26<br>(0.39;<br>23.50-25.02) | 14.36<br>(0.46;<br>13.45-15.26) | 13.89<br>(0.41;<br>13.09-14.69) | 13.89<br>(0.57;<br>12.78-15.01) | -3.15<br>(0.41;<br><0.0001) | -6.07<br>(0.36;<br><0.0001) | -10.37<br>(0.52;<br><0.0001) |

In comparing the difference in LS Mean for ATEC total score at Visit 1, all three pairwise comparisons between severity groups yielded statistically significant differences (Table 10). This is in contrast to Visit 8, at which point none of the comparisons reached statistical significance (Table 10).

**Table 10:** LS Mean differences between severity groups. Data are presented as: LS Mean difference (SE; P-Value)

|  | Visit 1 |  |  | Visit 8 |  |  |
| --- | --- | --- | --- | --- | --- | --- |
|  | Mild vs. Moderate | Mild vs. Severe | Moderate vs. Severe | Mild vs. Moderate | Mild vs. Severe | Moderate vs. Severe |
| **Total Score** | -6.50 (0.91;<br><0.0001) | -14.12 (1.55;<br><0.0001) | -7.62 (1.05;<br><0.0001) | 2.86 (1.27;<br>0.8449) | 4.20 (1.89;<br>0.8604) | 1.34 (1.49;<br>1.0000) |
| **Subscale 1: Communication** | -2.51 (0.23;<br><0.0001) | -5.46 (0.31;<br><0.0001) | -2.95 (0.26;<br><0.0001) | 0.46 (0.37;<br>0.9999) | 0.06 (0.47;<br>1.0000) | -0.39 (0.43;<br>1.0000) |
| **Subscale 2: Sociability** | -1.78 (0.22;<br><0.0001) | -3.51 (0.31;<br><0.0001) | -1.74 (0.24;<br><0.0001) | -0.50 (0.35;<br>0.9993) | 0.79 (0.45;<br>0.9887) | 0.29 (0.40;<br>1.0000) |
| **Subscale 3: Sensory** | -2.45 (0.33;<br><0.0001) | -6.75 (0.44;<br><0.0001) | -4.31 (0.38;<br><0.0001) | 0.46 (0.53;<br>1.0000) | 0.46 (0.68;<br>1.0000) | -0.01 (0.63;<br>1.0000) |
| **Subscale 4: Physical** | -5.24 (0.84;<br><0.0001) | -8.57 (1.17;<br><0.0001) | -3.33 (0.89;<br>0.0366) | -0.43 (1.31;<br>1.0000) | -4.55 (1.52;<br>0.2999) | -4.12 (1.33;<br>0.2341) |

The results for all four subscales mirrored those of ATEC total score, showing no statistically significant differences between severity groups at Visit 8 (Table 10). This may simply be an artifact of the definition of ASD severity, which is based solely on a child’s initial ATEC total score independent of child’s age. Therefore, a different approach to definition of ASD severity groups was investigated.

According to ATEC norms, ATEC total scores in young children decrease exponentially with age, with a time constant of approximately 3.3 years regardless of the initial ATEC total score (Mahapatra et al., 2018). Thus, participants could be divided into three approximately equal groups using exponents decaying with a time constant of 3.3 years. The exact parameters of exponents were determined using best-fit trendlines to ATEC norms (Mahapatra et al., 2018), Table 11).

**Table 11.** Severity group definition based on initial ATEC total score and age

| Severity group | Definition based on initial ATEC total score and age | Number of participants | % of participants in each group |
| --- | --- | --- | --- |
| Mild | $20 < \text{initial ATEC total score} \leq 17 + 119 * \exp(-\text{age}/3.3)$ | 710 | 31 |
| Moderate | $17 + 119 * \exp(-\text{age}/3.3) < \text{initial ATEC total score} \leq 27 + 189 * \exp(-\text{age}/3.3)$ | 805 | 35 |
| Severe | $27 + 189 * \exp(-\text{age}/3.3) < \text{initial ATEC total score}$ | 757 | 33 |

Group differences were reassessed using the severity group definition based on both the initial ATEC total score and age as specified in Table 11. The significance of interaction term between ASD severity group and Visits shows that the dynamics of ATEC total score and individual subscale scores varies between different severity groups (Table S35). Table 12 shows the LS Mean calculations for the three severity groups at Visit 1 and Visit 8. Reduction in ATEC total score was directly related to severity (Table 12). Over the two years the mild group improved by 16.46 units (SE= 0.92, p<0.0001), the moderate group improved by 21.27 units (SE=0.90, p<0 .0001), and the severe group improved by 20.48 units (SE= 0.90, p<0.0001).

**Table 12:** LS Means for various severity groups defined based on initial ATEC total score and age. Data are presented as LS Mean (SE; 95% CI). The difference between Visit 8 and Visit 1 is presented as LS Mean (SE; P-value).

|  | Visit 1 |  |  | Visit 8 |  |  | Visit 8 – Visit 1 |  |  |
| --- | --- | --- | --- | --- | --- | --- | --- | --- | --- |
|  | Mild | Moderate | Severe | Mild | Moderate | Severe | Mild | Moderate | Severe |
| **ATEC Total** | 57.58<br>(0.90;<br>55.80-59.35) | 62.82<br>(0.75;<br>61.35-64.29) | 66.15<br>(0.81;<br>64.55-67.74) | 41.12<br>(1.15;<br>38.87-43.36) | 41.55<br>(1.03;<br>39.52-43.57) | 45.67<br>(1.06;<br>43.58-47.75) | -16.46<br>(0.92;<br><0.0001) | -21.27<br>(0.90;<br><0.0001) | -20.48<br>(0.90;<br><0.0001) |
| **Subscale 1: Communication** | 15.22<br>(0.20;<br>14.83-15.60) | 15.41<br>(0.18;<br>15.06-15.76) | 14.94<br>(0.18;<br>14.60-15.29) | 9.42<br>(0.26;<br>8.92-9.92) | 10.60<br>(0.24;<br>10.12-11.08) | 11.90<br>(0.24;<br>11.44-12.37) | -5.80<br>(0.21;<br><0.0001) | -4.81<br>(0.21;<br><0.0001) | -3.04<br>(0.21;<br><0.0001) |
| **Subscale 2: Sociability** | 11.34<br>(0.25;<br>10.85-11.82) | 13.35<br>(0.22;<br>12.92-13.78) | 14.73<br>(0.23;<br>14.29-15.18) | 8.32<br>(0.33;<br>7.66-8.97) | 8.28<br>(0.32;<br>7.66-8.91) | 9.65 (0.32;<br>9.03-10.27) | -3.02<br>(0.30;<br><0.0001) | -5.07<br>(0.29;<br><0.0001) | -5.09<br>(0.29;<br><0.0001) |
| **Subscale 3: Sensory** | 12.97<br>(0.24;<br>12.50-13.45) | 14.37<br>(0.21;<br>13.95-14.79) | 15.19<br>(0.22;<br>14.76-15.62) | 9.23<br>(0.32;<br>8.61-9.86) | 9.30<br>(0.30;<br>8.71-9.89) | 10.29<br>(0.30; 9.71-10.88) | -3.74<br>(0.27;<br><0.0001) | -5.07<br>(0.27;<br><0.0001) | -4.90<br>(0.26;<br><0.0001) |
| **Subscale 4: Physical** | 17.47<br>(0.36;<br>16.76-18.18) | 19.86<br>(0.33;<br>19.22-20.50) | 22.52<br>(0.34;<br>21.87-23.18) | 13.50<br>(0.49;<br>12.53-14.45) | 13.31<br>(0.47;<br>12.42-14.25) | 14.92<br>(0.46;<br>14.01-15.83) | -3.97<br>(0.43;<br><0.0001) | -6.53<br>(0.42;<br><0.0001) | -7.60<br>(0.42;<br><0.0001) |

In comparing the difference in LS Mean for ATEC total score at Visit 1, all three pairwise comparisons between severity groups yielded statistically significant differences (Table 13). This is in contrast to Visit 8, at which point none of the comparisons reached statistical significance. For the Communication subscale all pairwise group differences were statistically significant at Visit 8, confirming the advantage of severity group assignment based on both initial ATEC total score and age and indicating that the mild group improved more than the moderate group and the moderate group improved more than the severe group (Table 13). There were no statistically significant differences between severity groups at Visit 8 in the Sociability, Sensory, or the Physical subscales.

**Table 13:** LS Mean differences between severity groups defined based on initial ATEC total score and age. Data are presented as LS Mean difference (SE; P-Value)

|  | Visit 1 |  |  | Visit 8 |  |  |
| --- | --- | --- | --- | --- | --- | --- |
|  | Mild vs. Moderate | Mild vs. Severe | Moderate vs. Severe | Mild vs. Moderate | Mild vs. Severe | Moderate vs. Severe |
| **Total Score** | -5.24 (0.84;<br><0.0001) | -8.57 (1.17;<br><0.0001) | -3.33 (0.89;<br>0.0366) | -0.43 (1.31;<br>1.0000) | -4.55 (1.52;<br>0.2999) | -0.41 (1.33;<br>0.2341) |
| **Subscale 1: Communication** | -0.19 (0.18;<br>1.0000) | 0.27 (0.22;<br>0.9999) | 0.47 (0.19;<br>0.7170) | -1.18 (0.30;<br>0.0144) | -2.48 (0.31;<br><0.0001) | -1.30 (0.30;<br>0.0029) |
| **Subscale 2: Sociability** | -2.01 (0.24;<br><0.0001) | -3.39 (0.30;<br><0.0001) | -1.38 (0.25;<br><0.0001) | 0.03 (0.40;<br>1.0000) | -1.33 (0.42;<br>0.2493) | -1.36 (0.40;<br>0.1106) |
| **Subscale 3: Sensory** | -1.40 (0.23;<br><0.0001) | -2.22 (0.29;<br><0.0001) | -0.82 (0.24;<br>0.0928) | -0.07 (0.38;<br>1.0000) | -1.06 (0.41;<br>0.6066) | -0.99 (0.38;<br>0.5689) |
| **Subscale 4:**<br>**Physical** | -2.39 (0.35;<br><0.0001) | -5.06 (0.43;<br><0.0001) | -2.66 (0.37;<br><0.0001) | 0.17 (0.58;<br>1.0000) | -1.42; 0.63;<br>0.8286) | -1.59 (0.59;<br>0.5206) |

## Discussion

The regular assessment of temporal change in symptoms of children with Autism Spectrum Disorder (ASD) participating in a clinical trial has been a long-standing challenge. A common hurdle in these efforts is the availability of trained technicians needed to conduct rigorous and consistent assessment of children at multiple time points. If parents could administer regular psychometric evaluations of their children, then the cost of clinical trials will be reduced, enabling longer clinical trials with the larger number of subjects.

The Autism Treatment Evaluation Checklist (ATEC) was developed to provide such a free and easily accessible method for caregivers to track the changes of ASD symptoms over time (Rimland & Edelson, 1999). Various studies have sought to confirm the validity and reliability of ATEC (Al Backer, 2016; Geier, Kern, & Geier, 2013; Jarusiewicz, 2002), yet none to date have assessed longitudinal changes in participants’ ATEC scores with respect to age, sex, and ASD severity. One trial conducted by Magiati et al., aimed to comprehensively assess ATEC’s ability to longitudinally measure changes in participant performance (Magiati et al., 2011). That study utilized ATEC to monitor the progress of 22 schoolchildren over a five-year period. ATEC score was compared to age-specific cognitive, language, and behavioral metrics such as the Wechsler Preschool and Primary Scale of Intelligence. The researchers noted ATEC’s high level of internal consistency as well as a high correlation with other standardized assessments used to measure the same capacities in children with ASD (Magiati et al., 2011). Charman et al. utilized ATEC amongst other measures to test the feasibility of tracking the longitudinal changes in children using caregiver-administered questionnaires and noted differential effects across subscales of ATEC, possibly driven by development-focused vs. symptom-focused subscales that are conflated in the ATEC total score (Charman, Howlin, Berry, & Prince, 2004). Another study assessing the ability of dietary intervention to affect ASD symptoms also utilized ATEC as a primary measure (Klaveness, Bigam, & Reichelt, 2013), concluding that it has “high general reliability” coupled with an ease of access. Whitehouse et al. used ATEC as a primary outcome measure for a randomized controlled trial of their iPad-based intervention for ASD named TOBY (Whitehouse et al., 2017). This trial was conducted over a 6-month time frame, with outcome assessments at the 3-month and 6-month time points. Although the study did not demonstrate significant ATEC score differences amongst test groups, the researchers reaffirmed their use of ATEC, noting its “internal consistency and adequate predictive validity” (Whitehouse et al., 2017). These studies support the viability of ATEC as a tool for longitudinal measurement of ASD severity which can be vital in tracking symptom changes during a trial.

The current study analyzed data reported by participants using the online version ATEC over a four-year time period from 2013 to 2017. Assessing these data permitted insight into the effects of age, sex, country of origin, and ASD severity on the longitudinal changes in ATEC score with all of these factors (save for sex) showing statistically significant differences affecting ATEC score dynamics. These findings identify specific variables capable of altering the developmental trajectory of children with ASD and indicate possible avenues of future investigation of causal relationships related to changes in ASD severity.

### Sex does not affect ATEC Score

The prevalence of ASD is strongly male-biased, affecting 4 times as many males as females. Accordingly, we were interested in differences in the rate of improvement between participants of different sexes. No significant differences in improvement of ATEC total score were observed. This suggests that the rate of improvement of ASD symptoms remains similar in males and females.

### Effect of Age on ATEC Score

The participants’ age was a significant modulating factor in determining the rate of their improvement. Younger children demonstrated greater improvement in ATEC total score. This phenomenon was recapitulated across subscales, with differences between the 2-3 YOA group and 3-6 YOA group reaching statistical significance for the Communication, Sociability, and Physical subscales and differences between the 2-3 YOA group and 6-12 YOA group reaching statistical significance for all subscales (Table 6). This finding is consistent with other ATEC longitudinal studies: younger children showed greater improvement in ATEC total score compared to the older children (Magiati et al., Charman et. al., Whitehouse et al., Table 14).

**Table 14.** Comparison of the annualized decrease of ATEC score across multiple studies.

|  | Number of<br>Participants | Age at<br>baseline<br>(years) | Initial<br>ATEC<br>total<br>score | Number of<br>ATEC<br>Evaluations<br>Completed by<br>each<br>Participant | Study<br>duration<br>(years) | Annualized change in score: LS Mean (SE) or mean±SD |  |  |  |  |
| --- | --- | --- | --- | --- | --- | --- | --- | --- | --- | --- |
|  |  |  |  |  |  | Total Score | Subscale 1 | Subscale 2 | Subscale 3 | Subscale 4 |
| **Current Study: 2-3YOA** | **407** | **2.6±0.3** | **71±22** | **5.9±3.7** | **2** | **-14.2 (0.6)** | **-4.3 (0.1)** | **-3.5 (0.2)** | **-3.2 (0.2)** | **-4.3 (0.1)** |
| Whitehouse et al./ TOBY<sup>A</sup> | 36 | 3.3±0.7 | 71±23 | 3 | 0.5 | -25.6±49.6 | -8.6 ±19.0 | 6.7±15.0 | -4.8±19.4 | -5.5±24.0 |
| **Current study: 3.1-6YOA** | **1205** | **4.3±0.9** | **60±24** | **5.4±3.5** | **2** | **-9.9 (0.4)** | **-2.4 (0.1)** | **-2.1 (0.1)** | **-2.3 (0.1)** | **-2.4 (0.1)** |
| Charman et. al. | 57 | 4.7±0.7 | 81±18 | 2 | 1 | -8.5±6.9 | -3.9±4.1 | -1.3±5.4 | -2.2±5.1 | -1.3±8.4 |
| Magiati et. al. | 22 | 5.6±0.6 | 53±21 | 2 | 4.8 | 1.3±3.6 | -0.5±0.6 | 0.7±1.3 | 0.3±1.1 | 0.9±1.8 |
| **Current Study: 6.1-12YOA** | **660** | **8.2±1.6** | **61±24** | **5.1±2.9** | **2** | **-6.9 (0.5)** | **-1.0 (0.1)** | **-1.6 (0.2)** | **-1.8 (0.2)** | **-1.0 (0.1)** |

<sup>A</sup> Includes both test and control groups as there was no difference in ATEC score change between the groups.

The magnitude of the annual decrease of the ATEC score was also found to be roughly similar to other reports across the studied age range. For the younger children the reduction of ATEC score seen in this study is in between those of Whitehouse et al./TOBY trial and Charman et al, Table 14. For the older children, the reduction of ATEC seen in this study is somewhat similar to that reported by Charman et al., Table 14.

The small differences between the studies can be attributed to differences in study design. In particular, the current study (1) had significantly more participants, (2) was based on greater number of ATEC evaluations, and (3) was conducted over the longer period of time than all the others discussed herein.

### Effect of ASD Severity on ATEC Score

In comparing the difference in LS Mean for ATEC total score at Visit 1, all three pairwise comparisons between severity groups yielded statistically significant differences (Table 10). This is in contrast to Visit 8, at which point none of the comparisons reached statistical significance (Table 10).

The results for all four subscales mirrored those of ATEC total score, showing no statistically significant differences between severity groups at Visit 8 (Table 10). This may simply be an artifact of the definition of ASD severity, which is based on a child’s initial ATEC total score. This method groups children with the same initial ATEC total score together independent of age. Thus, children who score 80 on their initial evaluation at the age 10 are grouped together with children who score 80 on their initial evaluation at the age 2. According to ATEC norms (Mahapatra et al., 2018), these children will score 70 and 25 respectively at the age of 12, and therefore clearly belong to different severity groups. This inconsistency in definition of ASD severity solely based on the initial ATEC total score independent of age may explain the observation that none of the group comparisons reached statistical significance at Visit 8.

The definition of ASD severity groups based on two parameters – the initial ATEC total score and age – yielded somewhat superior results compared to defining ASD severity based solely on the initial ATEC total score. While both definition methods showed no statistically significant differences between severity groups at Visit 8 in ATEC total score (Tables 10 and 13), the former method showed statistically significant pairwise differences between all the groups at Visit 8 for the Communication subscale, indicating more improvement in children with milder ASD and confirming the advantage of severity group assignment based on both initial ATEC total score and age.

### Role of Country of Origin

Conventional wisdom may suggest that the increased access to resources, including government-provided therapy for ASD, should lead to greater improvements. English-speaking nations (the United States, Canada, the United Kingdom, Ireland, Australia, and New Zealand) lead the world in government spending on therapy for children with ASD (Ganz, 2007; Horlin, Falkmer, Parsons, Albrecht, & Falkmer, 2014; Paula, Fombonne, Gadia, Tuchman, & Rosanoff, 2011) and therefore would be expected to produce superior outcomes of ASD therapy. Surprisingly, a comparison of English-speaking nations to the non-English-speaking countries demonstrated greater improvements in ATEC total score as well as in each subscale within the non-English speaking nations (Table 8).

This observation runs contrary to conventional thought and underscores the consensus that there is a potential for improving the treatment of children with ASD in the developed world. While it is difficult to speculate on the reason for this disparity between developed English-speaking countries and non-English-speaking countries, it is notable that child treatment is more often outsourced in the developed English-speaking countries compared to more traditional societies where grandparents are more commonly available and mother is more likely to stay at home to personally take care of a child (Fetterolf, 2017). Other factors, such as differences in diet (Adams et al., 2018; Rubenstein et al., 2018), reliance on technology (Dunn et al., 2017; Grynszpan, Weiss, Perez-Diaz, & Gal, 2014; Lorah et al., 2013; Odom et al., 2015; Ploog, Scharf, Nelson, & Brooks, 2013) and prescription medications (Lemmon, Gregas, & Jeste, 2011) could also play a role.

### Limitations

Participant selection presents a novel challenge in a study focused on caregiver-administered assessments. In the selection of participants for inclusion in this study, a baseline of ASD diagnosis could not have been established as child’s diagnosis is not part of ATEC questionnaire. Thus, it is not impossible that some of the participants did not have ASD diagnosis altogether. E.g., parents of a neurotypical toddler worried for any reason about an ASD diagnosis could have decided to monitor toddler’s development with ATEC evaluations and thus inadvertently added their normally developing child to the ATEC collection. As neurotypical children develop faster, the presence of neurotypical children in the dataset would have artificially increased the magnitude of annual changes of ATEC scores, predominantly for younger participants.

It is unlikely though that there were many neurotypical participants in our database. First, ATEC is virtually unknown outside the autism community. Second, there is a little incentive for the parents of neurotypical children to complete *multiple* exhaustive ATEC questionnaires (unless one of the children was previously diagnosed with ASD). Third, as described in the methods section, to further limit the contribution from neurotypical children, participants possibly representing the neurotypical population were excluded: those with an initial ATEC total score of 20 or less (7% of all participants) and those who completed their first evaluation before the age of 2 (3% of remaining participants). Despite this effort, the reported data may over-approximate the magnitude of annual changes of ATEC scores, especially in the younger participants.

As noted by other groups (Whitehouse et al., 2017; Charman et al., 2004), the use of ATEC as a primary outcome measure has some inherent drawbacks. While the ATEC is capable of delineating incremental differences in ASD severity amongst participants, the variety of measures amongst its subscales fails to differentiate developmental-specific changes from symptom-specific ones. This aspect of the ATEC may introduce a confounding variable when participants are at different developmental stages and follow unique developmental trajectories during a study. To mitigate these effects, trial designs must accurately separate participants based on developmental stage. This is most often accomplished by using age as a proxy for developmental stage.

### Conclusions

This manuscript attempts to characterize the typical changes in ATEC score over time as a function of children age, sex, ASD severity, and country of origin in a large and diverse group of participants. In doing so, it lends support to the efficacy of caregiver-driven psychometric observation, which when applied at scale, may be a viable alternative to using licensed technicians to assess the children.

## Acknowledgments

We wish to thank Dr. Petr Ilyinskii for productive discussion and scrupulous editing of this manuscript and Arthur Faisman for help with the database.

## Author contributions

Andrey Vyshedskiy, Edward Khokhlovich and Stephen M. Edelson designed the study. Stephen M. Edelson acquired the data. Edward Khokhlovich, Samantha Martinez, Benjamin Kannel, and Andrey Vyshedskiy analyzed the data. Shreyas Mahapatra, Edward Khokhlovich, Stephen M. Edelson, and Andrey Vyshedskiy wrote the paper.

## Compliance with Ethical Standards

Using the Department of Health and Human Services regulations found at 45 CFR 46.101(b)(4), the Chesapeake Institutional Review Board (IRB) determined that this research project is exempt from IRB oversight.

## Conflicts of Interest

The authors declare no conflict of interest.

